# Control algorithms underlying the translational optomotor response

**DOI:** 10.1101/2022.03.16.484575

**Authors:** John G. Holman, Winnie W.K. Lai, Paul Pichler, Daniel Saska, Leon Lagnado, Christopher L. Buckley

## Abstract

The optomotor response (OMR) is central to the locomotory behavior in diverse animal species including insects, fish and mammals. Furthermore, the study of the OMR in larval zebrafish has become a key model system for investigating the neural basis of sensorimotor control. However, a comprehensive understanding of the underlying control algorithms is still outstanding. In fish it is often assumed that the OMR, by reducing average optic flow across the retina, serves to stabilize position with respect to the ground. Yet the degree to which this assumption is true, and how this could emerge from the intermittent burst dynamics of larval zebrafish swimming is unclear. Here, we combine detailed computational modeling with a new approach to free-swimming experiments, in which *feedback gain* - i.e., the degree of sensory feedback produced by swimming - is manipulated by varying the height of larval zebrafish above a moving stimulus. We develop an account of underlying feedback control mechanisms that describes not only bout initiation but also the control of swim speed during bouts. We observe that the degree to which fish stabilize their position is only partial and gain-dependent, suggesting that the OMR may not primarily function in fish to prevent drift. We find the speed profile during bouts follows a fixed temporal pattern independent of bout intensity, suggesting that bout termination and bout duration are not explicitly controlled. We also find that the reverse optic flow, experienced when the fish is swimming faster than the stimulus, plays a minimal role in control of the OMR despite carrying most of the sensory information about self-movement. These results shed new light on the underlying dynamics of the OMR in larval zebrafish and will be crucial for future work aimed at identifying the neural basis of this behavior.

**Author Summary:** In many animals vision is central to the control of locomotory behaviors. In particular, innate motor responses to optic flow allow flying animals to react to gusts of air and fish to changes of current. As fish are washed downstream, for example, movement of the riverbed image across the retina evokes forward movement in bouts (short periods of swimming followed by rest periods). It is typically assumed that this translational optomotor response stabilizes the fish’s position to prevent it drifting downstream. In larval zebrafish, this response has become a key model system for investigating the neural basis of sensorimotor behaviors in a vertebrate. Here, we combine behavioral experiments and computational modeling to elucidate the underlying control algorithms. We unpick the detailed relationship between visual stimuli and the initiation, termination and intensity of swim bouts. We find initiation and intensity are controlled separately, and that termination is not explicitly controlled. Surprisingly we also found that the degree of stabilization is only partial and varies systematically with height above ground, raising questions of the function of this response. These findings shed new light on the underlying dynamics which will be crucial for future work to identify the neural basis of this behavior.

## Introduction

The optomotor response (OMR) plays a central role in locomotor behavior in animal species as diverse as insects(1,2), fish (3,4) and mammals (5) including humans (6). In zebrafish larvae it is one of the earliest and most robust behaviors to develop (7). The small size and translucency of the larval zebrafish makes it possible to carry out *in-vivo* whole-brain imaging studies (7,8), and the study of the OMR in this species has recently become a key model system for investigating the neural basis of sensorimotor control in a vertebrate (9–12). However, in comparison with insects, many aspects of the OMR in fish - the behavior itself, its function and the control algorithms underlying it - have received less attention (13). As Kist and Portugues point out (14) “… the OMR, a paradigm that has been extensively used, is still not fully understood, and its comprehensive characterization will undoubtedly reveal further insights into the neuronal circuitry underlying behavior”. Indeed, as Krakauer et al. argue (15), in most cases the neural basis of a behavior cannot be properly characterized without detailed prior understanding of the behavior itself and its underlying algorithms.

The OMR in young zebrafish larvae is a relatively complex behavior with two components (7,13,16). The usual account is that the fish first responds to rotational components of the whole-field optic flow by turning to face into the water stream. Second, they respond to translational components of the optic flow, specifically rostral/caudal movements across the retina of the image of the ground below, by swimming forward in a series of swim bouts (17) thereby counteracting the movement of its body relative to the ground caused by drifting backwards with the stream. The work reported here focuses exclusively on this second component, the *translational OMR*.

The translational OMR is often characterized as a locomotor behavior that acts in a similar way to the optokinetic reflex (OKR), i.e., by trying to bring the optic flow to zero, stabilizing the image on the retina, and thus also the position of the animal relative to the environment (12). However, far from stabilizing the image, intermittent bout swimming during the translational component of the OMR in fact causes the image to oscillate across the retina, moving rapidly forward during the relatively brief swim bursts and more slowly in the opposite direction when the fish is at rest between bouts. In this respect it is quite unlike the OKR which, at least during the slow phase, does stabilize the visual image (18). The “entrenched belief” (13) that the function of the OMR is to stabilize position relative to the ground has also been challenged, and indeed it remains unclear whether the degree of any regulation it achieves is sufficient in general to prevent drift.

Translational whole-field optic flow is certainly an important component of the effective stimulus for the translational OMR, but forward moving binary or steep dark-light transitions passing locally below the larva’s head also promotes triggering of swim bouts (14). Here we focus on the role of translational whole-field optic flow and use sinusoidal gratings that lack steep luminance gradients in order to maximize the relative importance of optic flow relative to other stimulus attributes.

Translational optic flow caused by linear movement relative to the ground is inversely proportional to the height of the eye above the ground (19). For flying insects, the importance of this factor has long been recognised. For example, maintenance of a near-constant optic flow by the translational OMR enables Drosophila to maintain a steady groundspeed that reduces steadily with height despite the perturbing effects of air currents (20) and honeybees to slow down and achieve a grazing landing as they approach the ground (21). For fish, changes in height while swimming are likely to be equally important, but have received much less explicit attention. For example, several studies (9,10,22) have varied *feedback gain*, defined in this context as the change in translational optic flow resulting from a given motor effort. Because translational optic flow is inversely proportional to height, so is the feedback gain. Although other factors such as fatigue, viscosity or temperature may also impact gain over longer timescales (10,22), for zebrafish larvae swimming in their natural habitat of shallow slowly moving streams in India it seems likely that changes in feedback gain will be dominated by substantial changes in height, e.g. halving when a fish rises to a position twice as far from the bottom, and that these will take place over much shorter timescales.

Optical imaging of neural activity normally requires a fixed-head procedure and measurement of a variable such as motor nerve activity (10,22) or tail movement (9) that is only correlated with swim speed; these variables do not provide a direct measure of swim speed and therefore of the absolute degree of regulation achieved by the OMR. In contrast, while free-swimming studies, like imaging studies, use a moving stimulus projected below the fish (23) to simulate the effect of the water current it is also possible to measure the actual swim speed. In at least one such free-swimming study (24) a close match between stimulus speed and mean swim speed consistent with good regulation has been observed for moderate stimulus speeds of up to about 15 mm/s. However, this experiment did not explore whether such a match is maintained over a range of different heights with different associated feedback gains.

Markov et al (12) proposed perhaps the most detailed and complete account to date of the translational OMR in larval zebrafish. It includes a model of a feedback controller that generates the behavior itself and proposes an additional mechanism that modulates parameters of that controller over longer timescales than those considered here to adapt to environmental changes. However their model makes the simplifying assumption that the feedback controller acts only to initiate and terminate bouts with swim speed during all bouts fixed at 20mm/s. In reality variation in mean swim speeds is correlated not only with changes in the timing of bout initiation and termination but also with variation in swim speeds during bouts (24). This is a prominent feature of the OMR and a complete account must include algorithms that control the intensity of swimming during bouts.

In this paper, we address some of the gaps in understanding through free-swimming experiments that manipulate feedback gain through changes in the height of the fish above a moving stimulus below, allowing absolute measurements of the degree of regulation achieved, and through detailed computational modeling. Together these lead to a number of new conclusions about the OMR and its underlying control algorithms. The experiments show that the degree of regulation achieved by the translational OMR is far from perfect and varies substantially, and systematically, with height and thus with gain. This casts doubt on the claim that its primary function is to maintain position. We also observe that the relative speed profile of individual swim bouts takes an invariant form that is essentially independent of bout intensity whether measured by initial speed, peak speed or total displacement. This suggests that although bout intensity and initiation are both explicitly controlled, bout termination and therefore bout duration are not.

Next, we present a series of algorithmic-level models of the OMR, based on the experimental findings and drawing initial inspiration from control theory, that propose mechanisms for the control of bout speed as well as bout initiation. First, we describe a basic model that proposes a single process of sensory integration implicated in both bout initiation and the control of bout intensity. This model proves sufficient to explain overall mean swim speeds but not the observed bout patterns. We then describe enhanced models in which optic flow is processed differently for bout initiation and intensity control and find that such a model can predict the bout patterns observed as well as mean swim speeds. We conclude that the control of bout initiation and bout intensity are at least partially decoupled: the control of bout intensity requires sensory integration while bout initiation appears to depend on the immediate unintegrated optic flow perhaps in conjunction with inhibitory motor integration. Furthermore, the final model suggests that although rostral translational flow induced by swimming carries much of the available sensory information about self-movement, surprisingly it may play little or no part in the control of either bout initiation or bout intensity and therefore of the translational OMR. Finally, we discuss some of the implications of these new findings.

## Results

### Free swimming experimental setup

We built a free-swimming behavioral rig (fig 1A) which mimics the situation where the larval zebrafish swim at various heights against a water stream by displaying visual motion below to induce translational optic flow. We tracked the 2D position of individual fish from above in separate longitudinal acrylic channels where sufficient length was provided for each of them to reach steady-state swimming for at least 90% of the trial distance. Moving gratings, scaled to ensure the spatial frequency remains constant and optimal (25), were projected on a computer monitor from below. The two independent variables, stimulus speed and height, were controlled via a custom-written program and manipulated through adjusting the vertical distance of the screen from the channels. We took a high-throughput approach and recorded batches of eight fish simultaneously, each in a separate channel. Using a mixed design to reduce disengagement from fatigue, each batch was subjected to only one of the testing height conditions but all the testing speed conditions.

**Fig 1.**
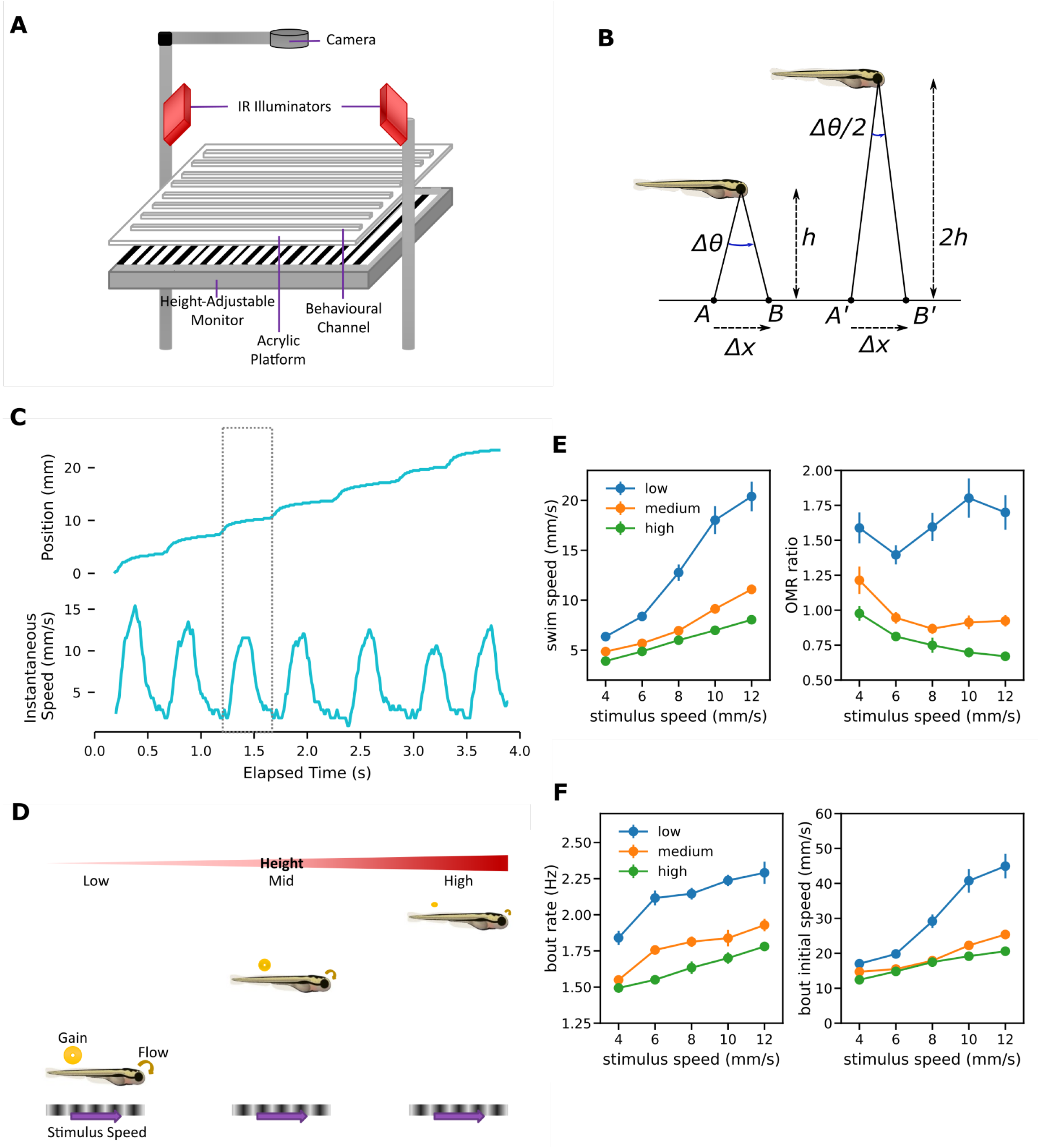
Free swimming setup and OMR regulation. (A) Schematic illustration of the experimental apparatus. (B) Optic flow is inversely proportional to height. The fish on the left is at rest a distance h above a moving stimulus. A point on the stimulus moves forward a small distance Δx from A to B over time Δt, causing the angle of the ray passing from that point through the optical center of the eye to change by Δθ with corresponding optic flow of Δθ/Δt radians/s. The fish on the right at twice a height experiences half the change in angle, Δθ/2, for the same stimulus movement in the same time, and thus half the optic flow. (C) Sample trace showing position of the fish from the start of the channel (top) and swim speed (below). The speed trace is derived from the position data and smoothed here for presentation. (D) OMR regulation procedure. Each fish is tested at one of three different heights and the same set of stimulus speeds. For a given stimulus speed, baseline optic flow - the optic flow experienced when the fish is at rest - and feedback gain both decrease as the height increases. (E) Mean swim speed and OMR ratio for the OMR regulation procedure. Error bars here and elsewhere: standard error (F) Bout patterns for the OMR regulation procedure.

### OMR regulation

During the translational component of the OMR, fish swim while maintaining an orientation that faces into the water stream thus reducing their speed over the ground. To examine the degree to which regulation is maintained despite the changes to optic flow induced by changes in height (fig 1B), and thus throw light on the underlying control mechanisms, we compared the swim speeds and degrees of regulation using the basic *OMR regulation procedure* in which different groups of fish swam at different heights with the grid below moving at the same set of fixed speeds (fig 1D). *OMR ratio*, defined as the ratio of mean swim speed to stimulus speed, served as the measure of regulation; perfect regulation corresponds to an OMR ratio of 1, under-compensation to a ratio less than 1 and over-compensation to a ratio of more than 1.

#### The degree of regulation varies systematically with height

Mean swim speeds increased with stimulus speeds (fig 1E). This is consistent with a regulatory effect, but the degree of regulation varied with height. Welch F tests showed a significant effect of height on both mean swim speed (F(2,56.1)=67.4, p<0.001) and OMR ratio (F(2,57.2)=70.9, p<0.001). At the intermediate height, fish swam at a similar speed to the stimulus. However, regulation at other heights was far from perfect: fish swimming at low heights overcompensated, swimming faster than the stimulus, while at a higher level tended to undercompensate. The overall mean OMR ratio for the intermediate level group, 0.97, did not differ significantly from the value of 1 associated with perfect regulation (t(31)=-0.62, p=0.54). However, the mean OMR ratio for the low level group, 1.62, was greater than 1 (t(31)=9.67, p<0.001) ; that for the high level group, 0.78, was less than 1 (t(31)=-7.57, p<0.001).

The increases in swim speed as height reduced resulted from increases in both bout rate and bout intensity (fig 1F). Bout rate was defined as the frequency of bouts occurring during OMR swim trajectories; bout intensity as the initial bout speed (the mean speed over the first 100ms). Welch F tests showed a significant effect of height on both overall mean bout rate (F(2,60.8)=82.2, p<0.001) and overall mean bout initial speed (F(2,57.8)=35.1, p<0.001)

### Feedback gain and the baseline optic flow procedure

*Baseline optic flow* can be defined as the optic flow experienced when the fish is not swimming, induced in the natural situation as the water stream sweeps the fish backward and in free swimming experimental procedures by the fixed forward movement of the grid stimulus. As feedback gain and baseline optic flow are both inversely proportional to height, the effects of each are confounded in the OMR regulation procedure.

In a free swimming setting the baseline flow can be held constant by increasing the stimulus speed in proportion to height. The resulting *baseline optic flow procedure* (fig 2A) enables feedback gain to be manipulated by changing the height while holding other factors constant, enabling investigation of the effects of behavior-induced sensory changes in an otherwise unchanged sensory environment.

**Fig 2.**
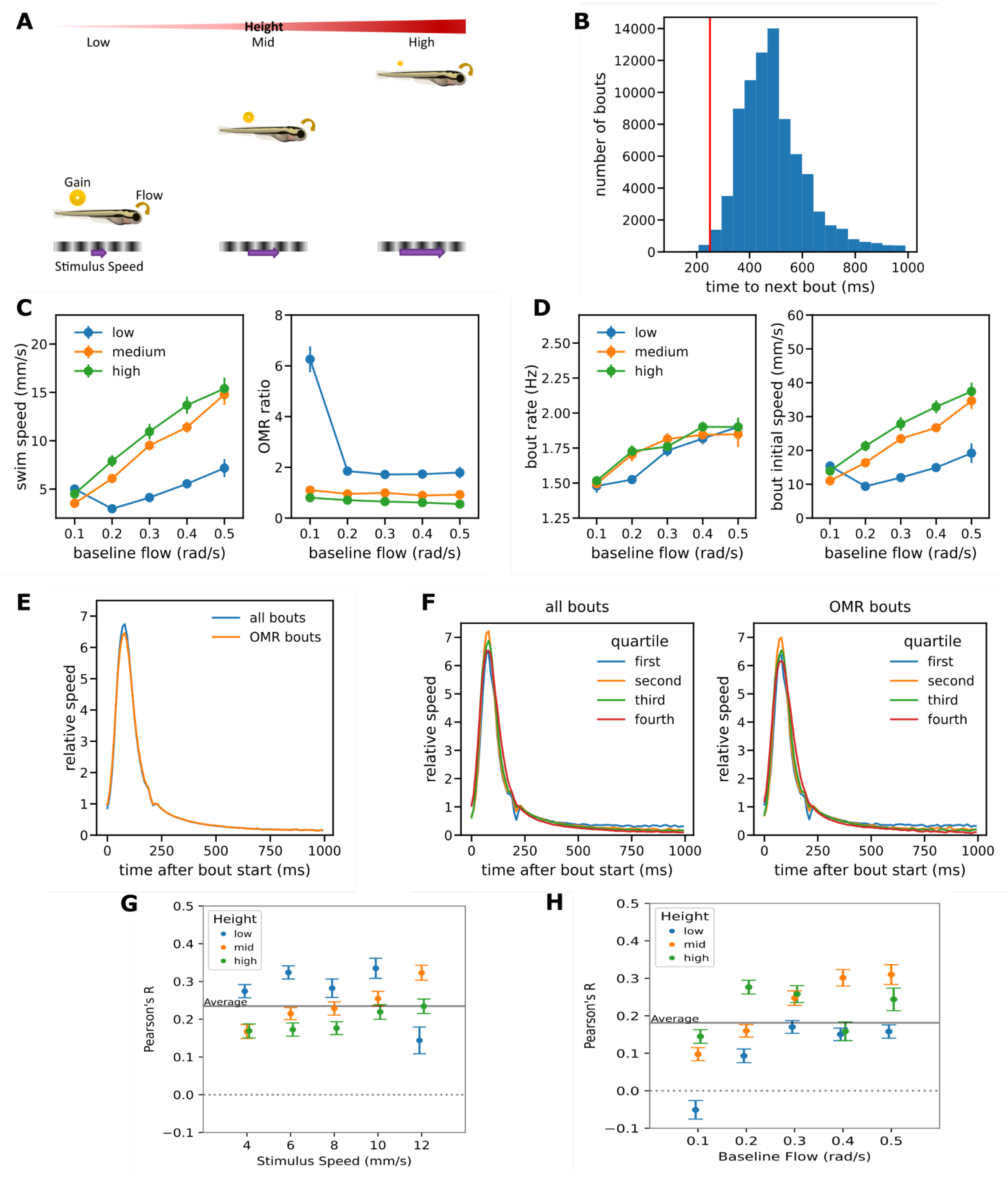
Experimental results for baseline flow experiment and bout analysis. (A) Baseline optical flow procedure. Each fish is tested at one of three different heights with the stimulus speed increased in proportion to height. Feedback gain decreases as the height increases but the baseline optical flow remains constant. (B) Histogram of inter-bout intervals for OMR bouts pooled from both procedures. The interval exceeded 250ms for 99% of bouts C) Mean swim speed and OMR ratio for the baseline flow procedure. (D) Bout patterns for the baseline flow procedure. (E) Overall bout speed profile averaged over both experiments for all observed bouts and OMR-consistent bouts only. The profile was not sensitive to the behavioral context. (F) Separate bout profiles for bouts split into quartiles by initial bout speed (swim speed averaged over the first 100ms of a bout). The relative speed profiles were essentially independent of bout intensity and again similar for all bouts (left) and OMR-consistent bouts (right). (G-H) Bout intensity and time to next bout are correlated. Correlation between bout intensity and time to start of the next bout for each experimental condition for the OMR regulation procedure (G) and the baseline flow procedure (H). In each case apart from the condition with slowest stimulus speed in the baseline flow procedure, for which behavior was generally anomalous, a positive correlation was observed.

This procedure is analogous to that of head-fixed virtual reality experiments in which feedback gain is manipulated by directly changing the relationship between swimming effort and stimulus speed while holding constant baseline stimulus speed and height, and therefore also baseline flow. Such experiments have shown that a variable correlated with swim speed such as the amplitude of tail movements (9) or motor nerve activity (2) typically increases as gain decreases even though at time points when swim bouts are initiated the environment remains constant.

Using this baseline flow procedure, we found that mean swim speeds increased with height even when the baseline flow is held constant (fig 2C), as expected from the virtual reality results (9,26). The sole exception was the condition with the lowest stimulus speed (0.8mm/s) which resulted in a very high OMR ratio of 6.26. This outlying data point was excluded from all further analyses as it seems likely that the very slow stimulus speed was insufficient to elicit the OMR, allowing instead the emergence of spontaneous swimming. As in the OMR regulation procedure, Welch F tests showed a significant effect of height on both mean swim speed (F(2,59.2)=50.9, p<0.001) and OMR ratio (F(2,55.7)=59.5, p<0.001).

To achieve full regulation in this experiment swim speeds for a given baseline optic flow must increase in proportion to height. The actual height above the stimulus for a given height condition could vary by up to 8mm (the height of the swimming channel), but the ratio of heights between the high- and low-level conditions was always at least 4.33. The largest observed ratio of swim speeds was only 2.67, showing again that the OMR achieves only partial regulation.

At times immediately prior to bout initiation the observed swim speeds were almost always very close to zero and thus optic flows close to the baseline level, suggesting that a reflex-like control mechanism based exclusively on the immediate optic flow at the point of bout initiation may not be sufficient. Instead, as others have argued (9,10), sensory integration is likely to be involved. However, because of the significant sensory delays associated with visual processing, when two bouts occur in quick succession a fish may still be experiencing some of the sensory consequences of the first bout, namely a reduction in the optic flow below the baseline level, at the time it initiates the second. The finding that swim speeds are sensitive to feedback gain as well as to baseline stimulus speed is therefore not sufficient to rule out a simple mechanism without further analysis.

#### Bout speed profile has a fixed form

To investigate the bout patterns underlying observed mean swim speeds we identified the times at which each bout was initiated. A bout was classified as *OMR-consistent* if it occurred when the fish was swimming in the same direction as the stimulus (see materials and methods section for details). Fig 2E shows the profile of instantaneous swim speeds for times up to 1s relative to the start of a bout averaged over all observed bouts, including those occurring when the stimulus was not moving, and normalized to give an average speed of 1 mm/s over that 1s period. The peak speed was reached about 80 ms after initiation and fell quickly thereafter to about 12.5% of the peak speed at 250 ms. A similar profile was observed when only OMR-consistent bouts were considered. The observed profiles also demonstrate that swim speed continues to decline from a low baseline for some time after active swimming has ceased; this is likely to be due to inertia.

Bouts were then split into quartiles by initial bout speed (defined as the mean swim speed over the first 100 ms) and separate profiles calculated for each quartile. These profiles were very similar to each other, whether considering all bouts or OMR-consistent bouts only (fig 2F), suggesting that the bout speed profile takes a fixed form that is insensitive both to the overall bout intensity and to the behavioral context (whether bouts occur as part of the OMR or another behavior such as exploration). Similar results were obtained when bout intensity was measured by peak speed (fig S1A) or total displacement (fig S1B).

The bout profile results imply that initial speed, peak speed and bout displacement are essentially equivalent as measures of bout intensity. We chose initial bout speed as it is less noisy than peak instantaneous speed and unlike bout displacement is not confounded with the occurrence of a closely following second bout.

#### Control of bout intensity and bout initiation are decoupled

The pattern of bout measures for the baseline flow procedure was qualitatively different from that for the OMR regulation procedure (fig 2D): changes in height still resulted in changes in bout intensity (F(2,59.8)=40.5, p<0.001) but bout rates remained essentially unchanged (F(2,59.9)=0.15, p=0.86). The finding that manipulating height affected mean swim speed via changes in both bout rate and intensity for the first procedure, but via bout intensity alone for the second, suggests that the mechanisms underlying bout initiation and intensity are at least partially decoupled.

### Modeling the translational OMR

#### General approach

Simple reflexes such as the tonic stretch reflex have often been modeled as feedback control systems (27). A similar approach may be taken for control of the OMR by treating optic flow as the feedback signal in a closed loop control system (12). In our formulation, an *optic flow processor* transforms the *sensed optic flow* to a *control signal* that drives a *bout generator* (fig 3A). The bout generator in turn outputs a motor signal reflecting the instantaneous swim effort as it varies through the bout, while the motor system is responsible for the detail of generating rapid tail beats that propel the fish forward through the water. The actual optic flow induced by the moving grid stimulus is proportional to the difference between the stimulus speed and speed the fish swims through the water, and inversely proportional to the height of the fish’s eye above stimulus (fig 3B; also see materials and methods). Here we assume that the sensory system provides an unbiased but delayed estimate of the actual translational optic flow.

**Fig 3.**
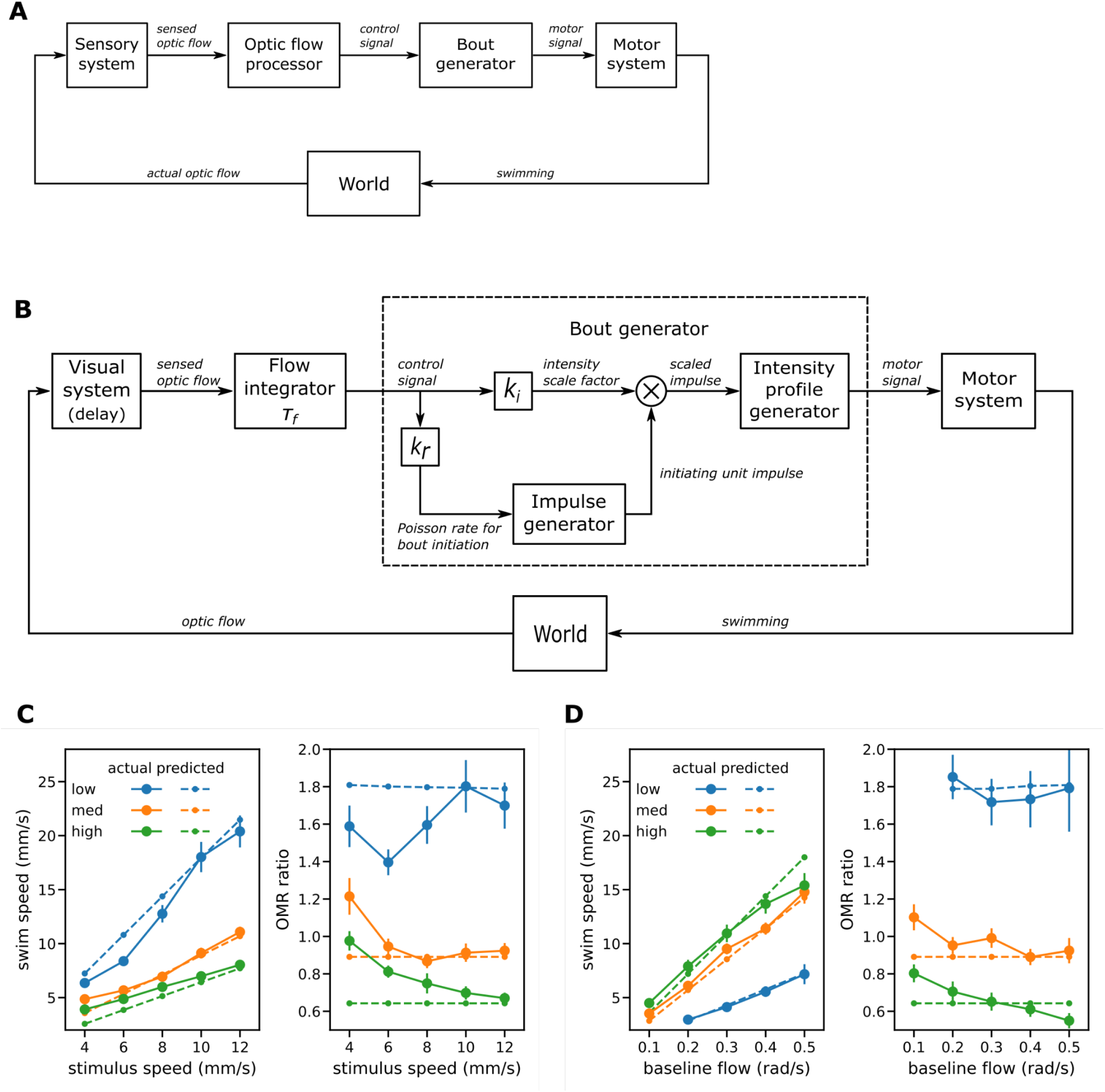
Modeling I: Single factor model. (A) Translational OMR as a feedback control system. (B) Block diagram for the single factor model. Optic flow processing consists of leaky integration of the sensed optic flow by the flow integrator. Within the bout generator, the integrated flow drives an impulse generator to determine when bouts begin and also modulates the amplitude of the resulting impulses input to the profile generator. The profile generator in turn generates the swim speed profile during bouts. (C-D) Comparison of swim speeds and degrees of regulation as observed in the experiments and predicted by the single factor model when fitted to observed swim speeds for the OMR regulation procedure (C) and the baseline flow procedure (D).

We present a relatively simple model that is able to predict mean swim speeds and therefore degrees of regulation. As might be expected given the observed decoupling of bout initiation and intensity, this model proved unable to predict detailed bout patterns, but serves as the basis of a series of more complex models presented subsequently.

From a control engineering perspective (28) we can view the optic flow processor as a controller and the combination of bout generator and motor system as the plant to be controlled. The usual control engineering task is to design a controller that achieves desired system behavior given knowledge of the plant; here the modeling task is to reverse engineer both the controller and plant to reproduce observed system behavior.

#### Optic flow processing and leaky integral control

Within engineering, most controllers found in practice are variants of proportional-integral-derivative (PID) control, a simple controller design that has been applied successfully to a vast range of control tasks (28). The input to such a controller is an error signal representing the difference between the desired and actual values of some observable variable. For proportional (P) control the output is just the immediate input error signal amplified by a gain factor, the *proportional gain*; for integral (I) control it is the integrated error signal amplified by the *integral gain*. For proportional-integral (PI) control a mixture of the two provides both timely response to changes in the error signal by virtue of the immediate proportional component and elimination of steady state errors by virtue of the integral component. For full PID control, the derivative of the error signal makes a further contribution, but this is very sensitive to sensor noise and often not used in practice (28); it is not considered further here.

When modeling the translational OMR it seems reasonable to identify the error signal directly with the forward-backward translational optic flow: to achieve regulation, if this component of the optic flow is positive (forward) on average the fish should speed up and if negative (backward) should slow down.

Traditional integral control, whether or not combined with proportional control, in general results in systems that have little or no steady state error. The experimental results above make clear that this is not the case for the OMR: significant degrees of over-compensation are observed for swimming at low level and under-compensation at high levels. This may not be surprising, as standard integral control requires perfect memory of the integral error while integration in biological systems is often considered to be leaky, as for example in leaky integrate-and-fire neuron models (29,30). The model for optic flow processing adopted here is therefore a generalization of standard integral control termed here *leaky integral (LI) control* in which the optic flow error signal is accumulated through time but also decays exponentially to zero with a fixed time constant (see Methods for details).

Leaky integral control could be combined with P control to create a generalized PI controller, but to keep the model as simple as possible we start by considering only LI control. However, P only control is included and treated as the limiting case of LI control as the time constant tends to zero.

#### Bout generation

A key difference between the swimming of zebrafish larvae and the output of most engineered systems is that movement is intermittent. The average swim speed and degree of regulation achieved depend on at least two factors: *bout intensity*, the swim speeds achieved during a bout, and *bout rate*.

Relative swim speeds during the bouts observed in these experiments followed a fixed temporal pattern (fig 2E-F) regardless of bout intensity. This observation suggests that swim bouts resemble fixed action patterns whose form is determined intrinsically rather than guided by external stimulation and that are ballistic in nature, i.e. once a bout is launched its speed trajectory follows a fixed course.

Markov et al (12) suggest that optic flow perception is subject to significant sensory delay of the order of hundreds of milliseconds; here we adopt their estimate of 220ms, which is similar to the duration of active swimming in the bouts observed here. As they point out, swimming cannot be affected by its impact on optic flow until after that delay period is over, implying that the initial portion of a bout is necessarily ballistic. The sensory delay rules out accounts of intensity profile generation based on visual sensory feedback; indeed, if their estimate of the delay is correct, the short bouts observed here must be ballistic for almost their entire duration as active swimming (defined as cessation of tail movements) was almost always less than 250ms.

These observations suggested modeling bout profile generation as a linear time-invariant (LTI) system whose impulse response is the observed normalized speed profile and whose input is a single short duration impulse at the time of bout initiation with magnitude proportional to bout intensity. This captures, in a simple way, the key features that bouts are ballistic with intensity following a fixed temporal pattern. One implication of such a model is that the swim speed trajectory of an individual bout is completely determined by the timing and magnitude of its initiating impulse; observed variations in bout duration (24) are secondary to variations in intensity and play no independent role in the control of swimming.

For bout initiation, we followed Portugues et al. (31) in proposing a stochastic process: bouts initiation events follow an inhomogeneous Poisson process whose instantaneous rate depends on the perceived optic flow. Another option is a noisy threshold model, but they found a Poisson process to give a better fit at least when modeling the latency for starting to swim following onset of stimulus movement.

Sensory delays in the visual system of the order of 220ms imply that the sensed optic flow remains high for almost the entire bout, as does the control input to the bout generator. To prevent immediate re-triggering of new bouts we therefore also imposed a *refractory period* during which bout initiation events are prohibited. The refractory period was not treated as a free model parameter but set from the experimental data at 250ms; the period between successive bouts was greater than this for over 99% of the bouts observed (fig 2B).

#### Single factor OMR model can predict mean swim speed

Figure 3B shows the simplest possible way to combine optic flow processing and bout generation as described above to yield a complete model of OMR control (see the methods section for mathematical details). A single control signal, the result of leaky integration of the sensed optic flow, governs both bout intensity and bout initiation.

The impulse generator implements the Poisson process and outputs a unit impulse when a bout initiation event occurs. This impulse is multiplied by the intensity scale factor to determine the scaled impulse input to the intensity profile generator, which in turn implements an LTI system whose impulse response is taken from the observed relative intensity profile. The motor signal output here thus reflects directly the swim speeds through the water to be achieved, which are then realized by the motor system.

Note that we did not attempt to model sensory or motor systems or the physics of the environment. A more complete model, for example, would specify the processes of motion detection including contributions from local features (14) and details of how tail beats are generated along with the physics of how the pattern of tail beats translates into propulsive force and how this translates into forward motion. Here however the focus is on the control of swim speed at the bout level, one level of abstraction above the tail beat level; we seek a model that given the environmental conditions of height and stream can predict characteristics of the bout patterns (rate and intensity) from which mean swim speed and therefore regulation emerges.

To fit the models, the data from both experiments were pooled and relative mean prediction errors calculated as described in the methods section. The parameter set that minimized the overall relative prediction error was then used to generate synthetic data from the model for both the OMR regulation and baseline flow procedures.

When only predicted mean swim speed is considered, optimizing parameters for the simple model to minimize swim speed prediction error alone gave a fairly good fit to the experimental data for both procedures (fig 3C, 3D). The main qualitative findings are replicated: in the OMR regulation procedures, fish swim faster than the stimulus when at a low level, at high level tend to swim slower, and at the intermediate height roughly keep pace (though the experimental data shows some deviations from this pattern at slow stimulus swim speeds).

#### The single factor OMR model does not predict bout characteristics

In contrast, the bout patterns generated by the model when fitted to mean swim speed differed greatly from those observed. In itself this is not surprising as the system is underdetermined: a given mean swim speed can be achieved by a high rate of weak bouts or a slow rate of strong ones. Refitting the basic model to minimize bout rate and bout intensity errors simultaneously rather than the mean swim speed error produced an improvement in bout characteristics, but the fit is still quite poor and predictions for mean swim speed are worse than when fitted directly to swim speed (fig 4A). In particular, the single factor model when fitted to the bout measures predicted that height has rather little effect on mean bout rates for the OMR regulation procedure but a large effect for the baseline flow procedure; the pattern observed was the opposite (fig 4C).

**Fig 4.**
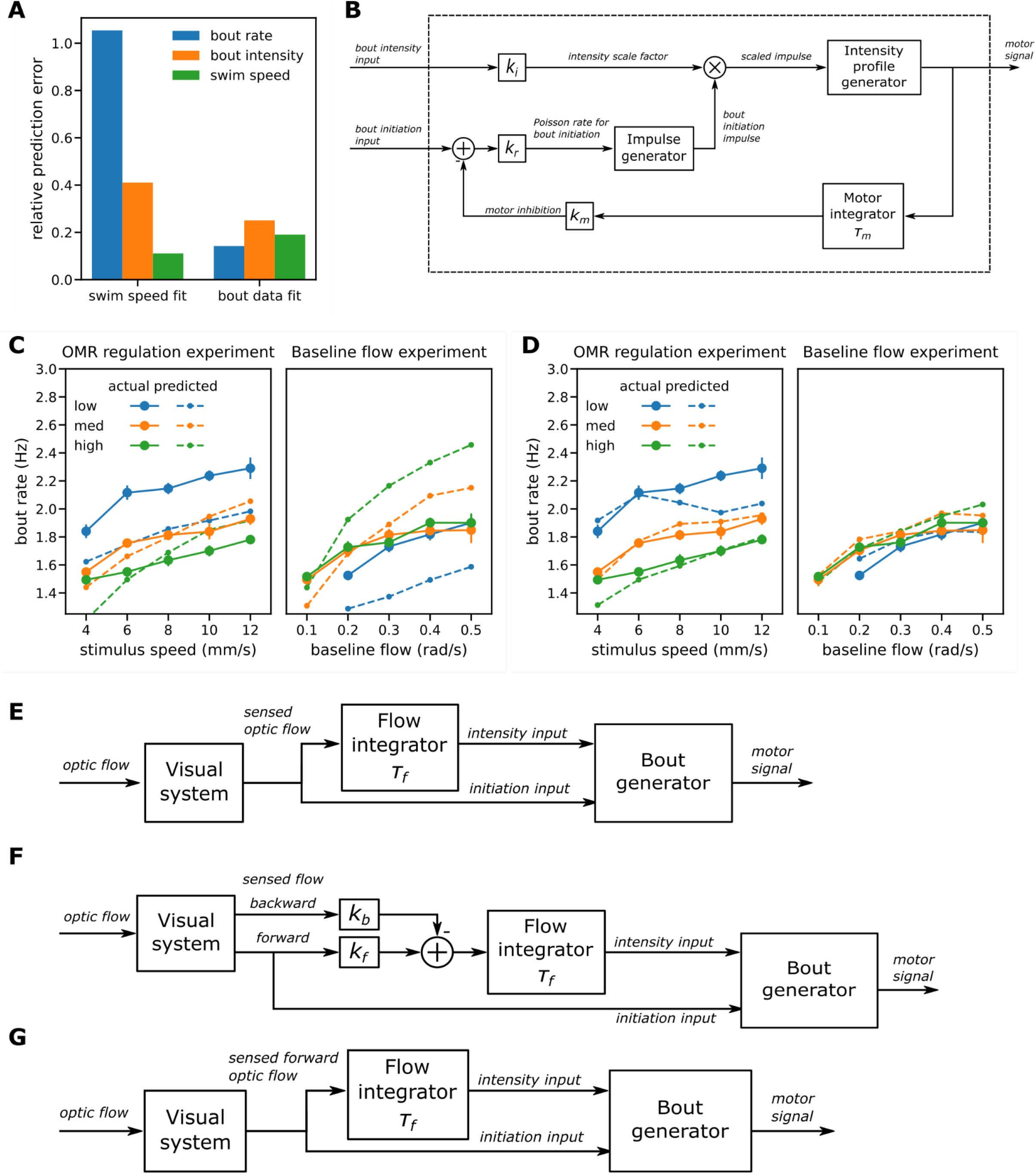
Modeling II: enhanced bout generation. (A) Mean prediction errors for the single factor model fitted to mean swim speed (left group of bars) and to bout measures (mean bout rate and intensity) (right group). Fitting to swim speed failed to predict the bout measures; fitting to bout data reduced prediction errors for bout measures but produced less accurate swim speed predictions. (B) Enhanced bout generator has separate inputs for bout intensity and bout initiation and the latter is subject to motor inhibition. (C) Bout rates for the single factor model fitted to bout measures. The model failed to predict that bout rates are sensitive to height in the OMR regulation procedure (left) but not the baseline flow one (right). (D) Bout rates predicted by the enhanced bout generator when fitted to bout data and constrained to reproduce the observed bout intensities. It was now possible to reproduce the bout rate patterns observed in the two experiments. (E-G) Dual factor model variant A. Variants A-C each have an enhanced bout generator with separate bout intensity and initiation inputs. Optic flow is integrated only for the intensity input. (E) Variant A: the overall optic flow is integrated. (F) Variant B: input to the flow integrator is a linear combination of the forward and backward components of optic flow. (G) Variant C: only the forward component of the optic flow participates in control of the translational OMR.

It seems then that the single factor model in which the same internal variable controls both bout rate and bout intensity can make reasonable predictions about mean swim speed but fails to capture how those speeds are realized at the bout level. This is not surprising: the baseline flow experiment suggested that bout intensity varies with height when baseline flow is controlled but bout rate does not, suggesting that the mechanisms responsible for bout initiation and for intensity are decoupled to a greater extent than can be accommodated by a single factor model.

#### Improving the bout initiation model

To investigate whether the bout generator component of the single factor model can reproduce the mean bout rates observed when relieved of the responsibility of also determining bout intensity, we set the intensity scale factor for the bout generator to be the mean of the bout intensities observed in the corresponding experimental condition and fitted the resulting model to the experimental data by minimizing the mean bout rate error alone. The relative bout rate prediction error was lower than before, but still substantial and the finding that bout rate is essentially independent of height when baseline flow is controlled still could not be reproduced (fig S2A). Importantly, when intensity was determined separately the best fit for bout rate was obtained with no integration at all (flow integrator output the same as its input) suggesting that bout initiation may depend only on the immediate visual stimulus.

What is needed for the model to replicate the finding that bout rate is independent of height when baseline flow is held constant? This is challenging if bout initiation is based on the visual stimulus alone. In the baseline flow experiment, fish swam during bouts more slowly at low level than at high level, but insufficiently so to compensate completely for the height difference. Consequently, the reduction from the same baseline in optic flow was substantially greater for fish swimming at low level than at high level. This might be expected to cause bout rate to reduce with height, as indeed is predicted by the model, but this was not observed in reality. Similarly reductions in height would act to increase the perceived gradient of local changes in contrast, and this although minimized here by the use of sinusoidal gratings would also be expected to increase bout rate (14).

In response we explored the possibility that an additional mechanism of motor inhibition participates in bout initiation. The basic idea is that the probability of a bout start event is reduced in relation to the recent amount of effort expended in swimming; in effect, all else being equal, stronger bouts are followed by longer recovery periods. The weaker bouts seen at low level in the baseline flow procedure result in less inhibition, counteracting the lower tendency to start swimming due to the greater reduction in flow sensed immediately following a swim bout and potentially resulting in similar mean bout rates at different heights.

Further analysis of the experimental data gave some support for this idea: for every experimental condition, apart from the lowest stimulus speed condition in the baseline flow procedure treated elsewhere as an outlier, there was a positive correlation between the strength of a bout and the time from the start of that bout to the start of the next (fig 2G-H)

The resulting bout generator model (fig 4B) has separate inputs controlling bout initiation and intensity, with motor inhibition only involved in the former: the motor signal is subject to leaky integration with time constant *τ*_*m*_ and gain *k*_*m*_ and the result subtracted from the bout initiation control input. In some respects this motor inhibition mechanism resembles that presented by Markov et al (12), but in their model bout initiation was based on a threshold mechanism rather than a Poisson process. Furthermore, they proposed a separate threshold for bout termination and did not model control of bout intensity: all bouts had the same fixed speed of 20 mm/s throughout and varied only in duration. In contrast, our account, because it does model swim speeds within bouts, has no need for a bout termination mechanism. Instead, following the fixed temporal profile, the swim speed reaches a peak and then decays to zero without any explicit termination event.

With the addition of motor inhibition, we found parameters that gave a better fit to bout rate (relative error reduced from 0.105 to 0.056) and replicated the finding that height has little impact on bout rate when the baseline flow is held constant. As for the basic bout generator the best fit was achieved when the flow integrator simply passes through the instantaneous optic flow signal without integration. This suggested that the model could be simplified by removing flow integration altogether from the signal path for bout initiation.

#### Dual factor model predicts bout characteristics as well as mean swim speed

Armed with the improved model for bout initiation we returned to developing a complete model that includes bout intensity. The bout initiation input of the enhanced bout generator (fig 4B) receives the sensed optic flow signal directly as it seems that flow integration is not required for that function. The intensity input receives an optic flow signal after leaky integration. For the initial dual factor model considered, variant A (fig 4E), the input to the flow integrator was assumed as before to be the overall sensed optic flow. Model parameters were again found that minimize the prediction error for mean bout rate and bout intensity simultaneously using the pooled data from both experiments. Variant A performed better than the basic single-factor model when fitted to bout statistics (fig 5A) but still failed to give a good match in some cases, in particular bout rate in the OMR regulation experiment (fig S3A)

**Fig 5.**
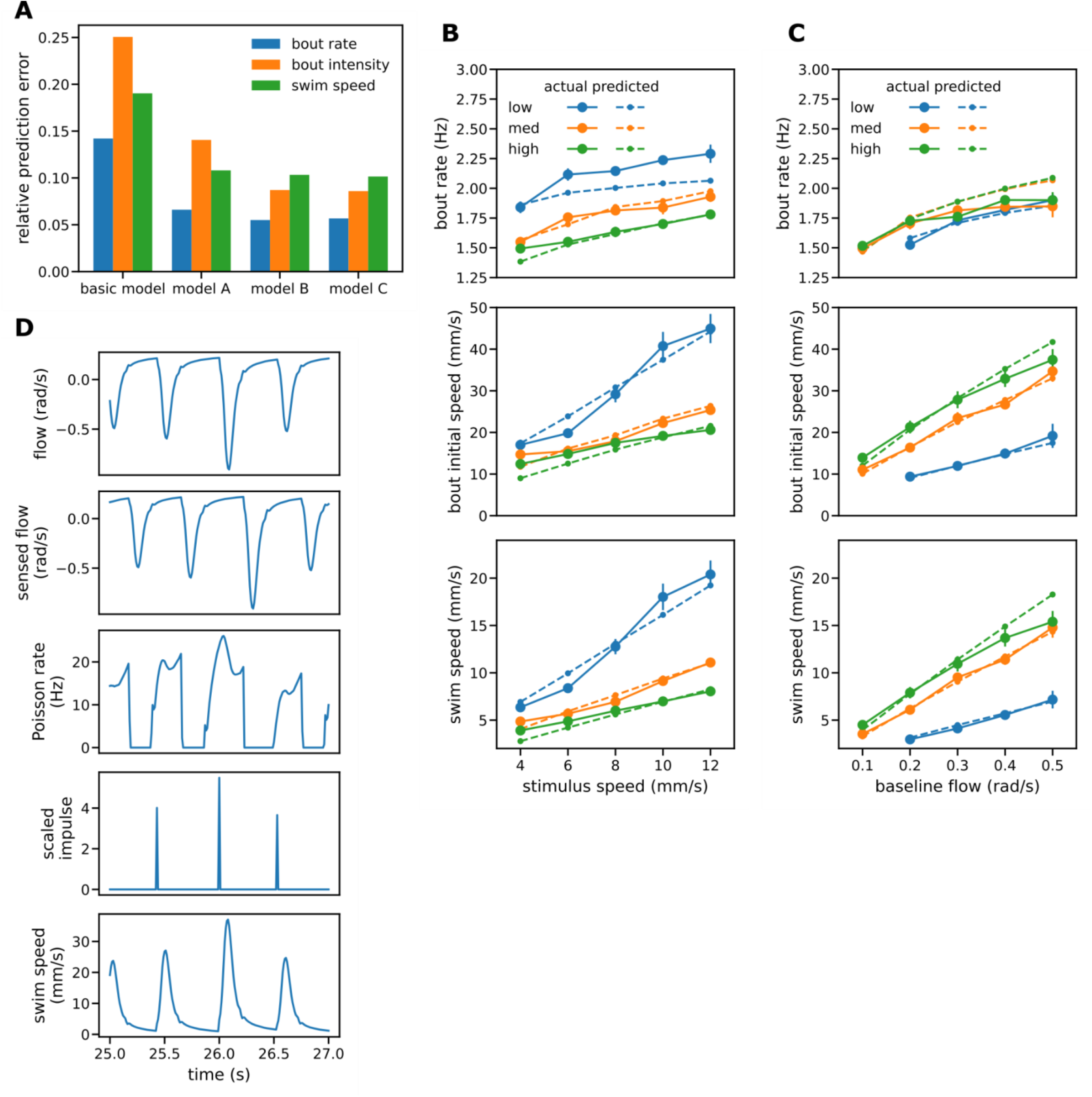
Modeling III: performance of enhanced model. (A) Relative prediction errors for the single factor model and three variants of the dual factor model. All variants of the dual factor model predicted both bout measures and swim speeds better than the single factor model. Variant C that processes only forward flow gave lower prediction errors than variant A and similar ones to the more general variant B, suggesting its adoption as the preferred model. (B-C) Comparison of observed bout rates, bout intensities and mean swim speeds with those predicted by variant C of the dual process model. In general, a good fit is achieved to the observed values for both OMR regulation (B) and the baseline flow procedures (C). (D) Sample signal traces generated by variant C of the two-factor model when height is intermediate (32mm) and stimulus speed 8 mm/s.

For variant A, the optimal time constant for flow integration turned out to be very low (0.024s) which might be expected to result in the model predicting a lower degree of regulation that is actually observed. A possible reason for finding such a low integration time constant is that otherwise the negative flow induced by swimming would result in an intensity control signal that stays negative for a significant amount of time after the bout finishes, preventing the timely generation of bouts with non-zero intensity. At any rate, such considerations suggested a model in which backward optic flow has a less powerful influence over bout intensity than forward flow.

To explore this we generalized the two factor model to associate different gains with the forward and backward translational flow components in a similar manner to Markov et al (12). Neurobiological evidence that different groups of neurons in the pretectum and tectum respond selectively to different directions of optic flow (32,33) suggest that such a scheme is biologically plausible. Input to the flow integrator is now a linear combination of the forward and backwards components parameterized by the different gains and the intensity gain parameter of the bout generator was fixed at 1 (fig 4F). For simplicity the bout rate input to the bout generator is shown as receiving the forward optic flow rather than the overall flow. This makes no difference to model output as without flow integration it is only the immediate optic flow that influences bout initiation and initiation events can never occur when the flow input is negative.

Allowing separate intensity gains for the forward and backward components of optical flow allowed a better fit to the experimental data (figs 5A, S3C, S3D). In addition, the best fit was achieved when the gain for backward flow was a very small proportion (0.027%) of that for forward flow, suggesting that backward optic flow has little or no influence on bout intensity as well as for bout initiation.

Accordingly, we simplified the enhanced model by setting the gain for backward optic flow to zero. For the resulting variant C (fig 4G) the input to the flow integrator is the forward component of optic flow rather than the overall flow; it is otherwise identical to the initial variant. Sample signal traces are shown in figure 5D. After optimizing parameters for variant C starting from the optimal parameters found for variant B, both variants gave similar prediction errors (fig 5A). The optimal time constant for flow integration for variants B and C was considerably greater than that for variant A, from 0.024s to 0.152s, supporting the idea that participation of backward optic flow reduces model fit for bout intensity as well as bout initiation. As model C is simpler than model B but gives similar results we selected it as the final model.

Comparison of observed bout rates, bout intensities and mean swim speeds with those predicted by the dual final model showed a generally good fit for all measures for both for the OMR regulation procedure (fig 5B) and baseline flow procedure (fig 5C). In particular the final model reproduced both qualitative experimental findings: that the degree of regulation varies systematically with height and that bout rate is relatively insensitive to height in the baseline flow procedure but not the OMR regulation procedure while bout intensity varies with height for both.

## Discussion

### Key features of the translational OMR

#### The degree of OMR regulation is only partial and varies systematically with height

In experiment 1, which simulates the natural situation in which a fish may swim at different levels against an approximately constant water stream, some degree of regulation was observed. However it was far from perfect, with mean swim speeds varying systematically with height: fish tended to over-compensate (swim faster than the stimulus) when close to the bottom and to under-compensate at higher levels.

#### Bout initiation and bout intensity are separately controlled

In experiment 2, the free-swimming analogue of VR procedures that vary feedback gain while holding the baseline stimulus speed constant, we found that mean bout intensity reduces as height reduces (gain increases) but mean bout rate remains unchanged. This dissociation suggests that different mechanisms underlie the control of bout initiation and bout intensity. Modeling reinforces that conclusion, suggesting that control of bout intensity depends on the leaky integration of forward optic flow while bout initiation is determined by the immediate forward optic flow in conjunction with an inhibitory effect of recent motor activity.

#### Bout speed and bout duration are not separately controlled

The temporal profile for relative swim speeds during swim bouts appears to be independent of bout intensity whether measured by initial bout speed, peak speed or total displacement. This invariance suggests that the absolute swim speeds observed during a bout can be characterized by a single value which acts as a scaling factor applied to a fixed relative speed profile. Correlations between bout speed and bout duration as reported in previous studies (24) might be expected with the details depending on how bout termination events are identified (e.g. swim speed or tail curvature falling below an absolute threshold), but an invariant relative profile suggests that bout speed and duration share a common causal factor and are not independently controlled. Moreover the relative speed profiles for OMR-consistent bouts were essentially identical to those observed when the stimulus was at rest or moving in the wrong direction, suggesting that the bout generation mechanisms described here are the same for all behavioral contexts.

#### Only forward optic flow plays a significant role

The results of modeling suggest that the effective external stimulus for the translational OMR is restricted to the forward component of the whole-field translational optic flow, with backward flow playing no significant role in the control of either bout initiation or bout intensity. This may seem surprising under the common assumption that the OMR serves to minimize drift, since the backward optic flow provides information that should make good regulation easier to achieve, but is consistent with the finding that regulation is generally quite poor.

### Characterizing the OMR

The optomotor response is sometimes called the “optomotor reflex” (5,34), though more often when referring to the rotational rather than the translational component, or characterized as a “position-stabilizing reflex” (11). However, the results presented here suggest that at least the translational component of the OMR is not a typical reflex. In a reflex action, “neural activity evoked by stimulation of sensory receptors is transmitted directly to response production mechanisms” (35). However, the translational component of the OMR, a behavior that continues over a sustained period of time, is not a direct response to an immediate eliciting stimulus but requires intermediate processes that involve integration and therefore a form of memory both of the optic flow for control of bout intensity and of motor activity for control of bout initiation.

It has also been suggested that the translational OMR is an adaptive locomotor behavior that involves a form of motor learning (9,10) but the current model suggests that its control, at least over relatively short time scales, does not require learning - at least if learning is defined as a persistent change in behavior brought about by exposure to specific environmental events during the organism’s lifetime. Similarly, as Markov et al argue (12), “adaptation”, which implies a persistent change in the brain in response to environmental or bodily changes, does not seem to be required for the moment-to-moment control of the OMR.

Instead, the OMR is probably an innate sensorimotor behavior that emerges through development. Swimming behavior during individual bouts in some ways resemble a traditional “fixed action pattern” (36) or “modal action pattern” (35,37) whose release and overall vigor depends on external stimulation while the detailed pattern of behavior takes a stereotyped form typical of the species that is generated intrinsically rather than determined by such stimulation. The OMR as a whole might then be characterized in ethological terms as a stream of modal action patterns of varying intensity although in this case the immediate releasing stimuli are internal rather than external and generated through processing of the whole-field forward optic flow and a motor-related signal.

### Neural mechanisms underlying the OMR

As an algorithmic level model (38) the current model has nothing directly to say about how component operations are implemented at the circuit level or map to different brain regions, but it does both constrain and suggest possibilities for an underlying neural implementation. For the latter, one possibility is that the the visual processing, including integration, is a function of pretectal regions, generation of bout initiating impulses a function of midbrain or hindbrain areas, and that a central pattern generator (CPG) located in the rostrocaudal spinal cord is responsible for the detailed pattern of tail movements. There is neurobiological evidence for the first (11,12,14,32,33) and last of these (16,39) but less evidence relevant to the proposed intermediate processes. Outstanding questions include how the magnitude of the logical impulse received by such a CPG is encoded at the neural level. For example each bout might be triggered by a brief burst of firing where some combination of duration, frequency, and/or the number of neurons involved corresponds to the magnitude of the scaled impulse in the algorithmic model. Another is the locus of motor integration. One possibility, as suggested by the block diagram for the bout generator, is that it is collocated with impulse generation, but it could equally be located in motor or even sensory areas.

### Function of the OMR

The prevailing account of the function of the OMR is that it serves to hold position relative to the ground (13). The finding here that regulation on average is far from perfect throws that hypothesis into doubt, but is not sufficient to refute it. For example, perhaps the OMR in 6-7 dpf larval zebrafish is still developing with regulation improving later in life, although the observation that older zebrafish larvae (16-18 dpf) and adults tend to exhibit a negative translational OMR (13) - i.e. swim along with the current rather than against it - makes that seem less likely. Another possibility is that good regulation requires, in addition to optical flow stimulation, local visual stimuli absent from a sinusoidal grating (14). Hydromechanical cues such as turbulence that are not available when testing in still water may also play a role as in moving water such cues can be detected by the lateral line system and in the absence of vision are sufficient to elicit a degree of counterflow swimming (40). However, at least in 5-9dpf larval zebrafish, once counterflow swimming has begun, the degree of regulation in the presence of visual cues seems to be the same whether or not lateral line stimulation is also available (40). A final possibility is that OMR behavior varies widely between individuals, with only a few achieving good regulation. The vast majority of individual larvae do not reach adulthood for many reasons perhaps including being swept downstream; those that do may be good regulators for whom the OMR achieved its assumed function of holding position.

Alternatively, the OMR may contribute to fitness for a different reason. One possibility is to coordinate the swimming of neighboring individuals: all will tend to swim in the same direction when subject to a similar water current. (There is some evidence for a similar effect of the rotational OMR in the adult medaka (41), though not in larvae). This may tend to keep individuals together reducing vulnerability to predation at a time when shoaling behavior has not yet developed. Other suggestions include energetic benefits of swimming into the stream and enhanced escape from suction predators (13,40). Finally, perhaps even partial regulation confers some benefit by reducing drift: swimming is required for individuals to come into proximity with food and the OMR promotes an optimal direction and some adaptation to the strength of water current. In fact there may anyway be little additional fitness benefit to an OMR that holds position perfectly when performed in isolation: in natural settings this behavior is likely to occur in alternation with prey capture and other behaviors, as observed in some insects (42), whose fast movements would disrupt regulation. Fieldwork and further experimentation would be required to evaluate these possibilities.

### Other model variants

The current model proposes a non-homogeneous Poisson process for generation of initiating impulses and that motor inhibition results from integration of a motor signal that reflects the observed swim speed. Other variants are of course possible. For example, with the involvement of motor inhibition a mechanism that generates a spike whenever the sensed optic flow first exceeds the inhibitory motor signal by some threshold might be sufficient. The inhibiting signal might result from leaky integration of the scaled impulse rather than a more downstream motor signal, giving a faster response that might even remove the need for a separate refractory period. Future work includes exploration of such options.

Modeling the input signal to the intensity profile generator as an impulse is the simplest possible approach as it allows the impulse response of the profile generator to be recovered directly from observed behavior. In reality, that is likely to be an oversimplification: as suggested above when considering the neural implementation, bout intensity may reflect the duration of an initiating pulse instead of or as well as its magnitude. Recovering the impulse response from observed behavior would then require deconvolution and knowledge or assumptions about the form of the input signal, including incorporation of relevant local features (14) as well as whole-field optic flow.

### Are swim bouts always ballistic?

Some other studies have been interpreted as suggesting that swim bouts are not always ballistic but may be modified by external stimulation once past the initial period of sensory delay. For example Markov et al (12) observed faster swimming after the initial 220ms of a bout in a VR situation for open loop bouts, when grid movement was unaffected by swimming, than for closed loop bouts. They suggested that bouts consist of an initial ballistic period followed by a subsequent reactive period. In contrast, the current model suggests that the original bout is entirely ballistic; instead, continued exposure to forward optic flow in the open loop condition triggers a second bout whose impact on swim speed merges with that of the first. Further research is required to decide which interpretation is correct.

## Methods

### Ethics Statement

All procedures were in accordance with the UK Animal Act 1986 and were approved by the Home Office and the University of Sussex Ethical Review Committee under the project license PPL70/8851.

### Data and code availability

All data and code for experimental data analysis, model simulation and fitting are available from a GitHub repository at https://github.com/johnholman/omr-algorithms

### Animals

Wild-type zebrafish (Danio rerio) adults were housed in the aquatic facility in the University of Sussex. All experiments were performed on larvae aged 6 to 7 days post-fertilization. Larvae were reared in Petri dishes in E2 solution on a 14/10 hour light/dark cycle at 28°C. Pebbles were added in Petri dishes and growing larvae were put above white noise grids for habitat enrichment and early visual feedback.

### Free-swimming behavioral assay & procedures

We used a free-swimming assay to mimic the natural environment in which larval zebrafish swim against aquatic perturbations. Larvae were put in 300mm long acrylic channels with depth 8mm. Moving sinusoidal gratings with 100% contrast were presented from below to simulate the optic flow experienced in streams. Grating speed was kept relatively low to avoid eliciting escape behaviors (2-12mm/s) and grating width varied to maintain the same optimal spatial frequency between 0.01-0.02 cycles per degree regardless of height (25). Height was varied by adjusting the stimulus monitor within the range of 4-60mm. Position of the fish was tracked through the centroid of the 2D image taken from above at 100 frames/s using Basler acA1920-155um USB 3.0 camera with a resolution of 2.3 MP. The setup was illuminated by IR light at a 45 degree angle from above on the side.

Eight larvae were tracked simultaneously in parallel channels and each experimental session consisted of three traverses at the same height. The initial traverse was to drive all larvae to one end of the channel with the stimulus moving at 5 mm/s for 30s. During the two trial traverses the grating moved first forward then backward at the appropriate speed for that session with a resting period of 5s between trials. Experiments were conducted on two consecutive days with larvae aged 6 and 7 dpf respectively. The monitor presenting the stimulus was manually adjusted for the three experimental heights at 4-12, 28-36 and 52-60 mm respectively. We took a mixed design approach: each batch of 32 larvae (total of 96 per experiment) was subjected to only one specific height but all the stimulus speeds. The order of experimental conditions with different combinations of testing height and speed was randomized on the day.

### Experimental data processing

#### Tracking algorithm

OpenCV blob detection algorithm was developed in Cython to handle the camera for recording the x and y positions of multiple fish simultaneously in every frame. Maximum light reflection from the swim bladder of each fish was identified as the center of mass. Data acquisition was done in real-time to output positional time-series in each channel as manually defined. Positional time-series data was then cleaned by a custom-written Python script using Markov chains to predict the actual trajectory of each sample from surrounding noise and substitute missing points.

#### OMR trajectory extraction

Position data from the tracking algorithm were first resampled with a fixed timestep of 10ms. Time gaps of up to 120 ms were filled by linear interpolation and the rest left as missing data.

An *OMR trajectory* was defined as a time series of duration at least 1s with no remaining missing values during which the mean swim speed was at least 0.5 mm/s and the direction of swimming the same as that of the stimulus. Times at which swimming changed direction were defined as zero crossing times for the difference between two exponential moving averages (EMAs) of position. The EMA at time t for positions *x*_0_ … *x*_*t*_ with smoothing factor α was calculated using the *ewm()* method in pandas (43) as

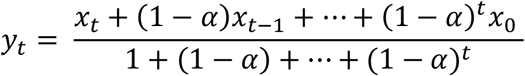

α was 0.002 for the slow decay EMA and 0.05 for the fast decay EMA. The direction of time was reversed when looking for the end point of a trajectory to ensure that swimming was in the appropriate direction throughout.

The resulting OMR trajectories were pruned to exclude timesteps following the first arrival at a point 50mm before the physical end of the channel for each of the two periods during which the stimulus was non-stationary.

#### Bout extraction

A low-pass bidirectional filter with a ‘kaiser’ window which has a length of 20 and cut-off frequency of 0.02 was applied to the position to smooth the data. Derivative functions were then used to obtain the velocity and acceleration. Bout onsets and offsets were defined as local maxima and minima and any unpaired point was discarded. Only bouts with a duration of more than 50ms and a distance of at least 0.5mm were considered valid.

### Statistical analysis

R (44) was used for statistical analysis. Welch F tests were used to assess the impact of height on the dependent variables (swim speed, OMR ratio, bout rate and bout intensity) as the requirement of homogenous variance for classical Fisher ANOVA was not met, with observations for the repeated measures variable (stimulus speed in experiment 1, baseline flow in experiment 2) averaged for each individual.

### Model description

#### Optic flow and sensory model

Movement of the fish relative to stimulus below generates an optic flow ω in radians/s:

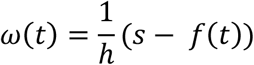

where *f*(*t*) is the speed of the fish through the water at time *t, s* the forward speed of the grid stimulus (this corresponds in the natural setting to the speed of the water current sweeping the fish backwards), and *h* the height of the fish above the water. The convention here is that optic flow is positive when the fish is at rest with the stimulus moving and is reduced by swimming. The output of the fish visual system, the sensed optic flow *y*(*t*), is assumed to be an accurate estimate of the actual optic flow but with a sensory delay *t*_*delay*_ of 220ms:

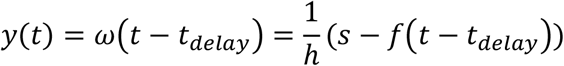

#### Leaky integration

Integration is assumed to be subject to a “leak” that causes output to decay exponentially to zero in the absence of input. Optic flow integration is modeled in continuous time by the ODE

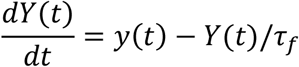

where *Y* is the output of the integrator and *τ*_*f*_ is the decay time constant. This is equivalent to a low pass IIR filter with time constant and gain both *τ*_*f*_; for a fixed input *y* the output *Y* relaxes exponentially toward asymptotic value *yτ*_*f*_.

For model simulation the discrete time approximation uses Euler integration with timestep number *t* and time step duration *Δt* of 10ms:

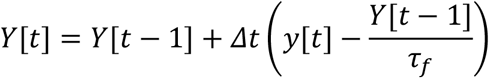

#### Initial model

For the initial model, the output of the flow integrator is the only input of the bout generator and governs both bout initiation and bout intensity. Within the bout generator, bout initiation events are represented by a train of unit impulses. These are emitted by an impulse generator implementing a nonhomogeneous Poisson process whose instantaneous rate is *max*(0, *k*_*r*_*Y*(*t*)) with *k*_*r*_ a fixed model parameter and output *z*(*t*). To avoid immediate re-triggering we also require a minimum period of 250ms, the refractory period, between consecutive impulses.

The intensity of a bout initiated at time *t*_0_ is determined by *k*_*i*_*Y*(*t*_0_) with *k*_*i*_ another fixed model parameter. The intensity profile generator is modeled as a linear time invariant (LTI) system. Its input is the train of scaled impulses *k*_*i*_*Y*(*t*)*s*(*t*) and its impulse response *g*(*t*) can be estimated directly from the experimental data input. Its output is

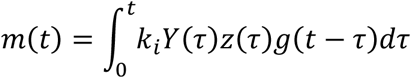

For the corresponding discrete time approximation for simulation of the impulse generator, the probability that a swim bout is initiated on timestep *t* is

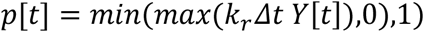

The profile generator LTI for simulation has impulse response *g*[*t*] and we simplify by assuming that speeds during a bout are independent of the history of previous bouts and thus determined only by the magnitude of its initiating impulse. The motor output is therefore simply

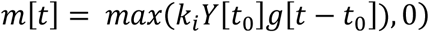

where *t*_0_ is the time step at which the most recent bout began.

The motor system translates the motor signal to swimming movements that generate a propulsive force resulting in movement through the water. Adopting the simplest possible physical model, we assume that inertia is negligible relative to viscous friction and that there is no delay associated with the motor system so that swim speed through the water *w* is proportional to the instantaneous propulsive force and that propulsive force is proportional to the motor signal. In addition, the model does not attempt to account for the effects on performance of variables such as fatigue, water temperature and water viscosity, factors that are held as constant as possible during the experiments. We therefore assume both proportionality coefficients are also constant and thus can be absorbed into the existing parameter *k*_*i*_ to identify swim speed *f* directly with motor output:

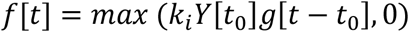

#### Two-factor model with motor inhibition

For the general two-factor model the visual system is assumed to provide separate equally delayed accurate estimates of forward and backward optic flow, *y*_*f*_(*t*) and *y*_*b*_(*t*), with the convention that both are positive so the overall sensed flow is *y*_*f*_(*t*) − *y*_*b*_(*t*). The input of the flow integrator is a linear combination of the two with positive coefficients *k*_*f*_ and *k*_*b*_ so the output of the flow integrator in the discrete-time simulation is

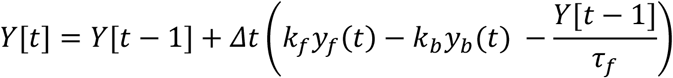

The associated bout generator (figure 3G) has two inputs. The first input is driven from the optic flow integrator output and determines bout intensity while the second is driven directly by the forward component of sensed optic flow and influences bout initiation. The generation of bout initiation impulses is inhibited by an additional motor feedback loop that reduces the instantaneous Poisson rate. For the corresponding discrete time approximation, the output of the motor integrator with leak time constant of *τ*_*m*_ is

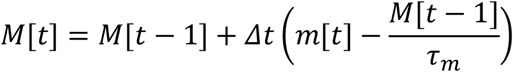

The probability of bout initiation during timestep *t* is

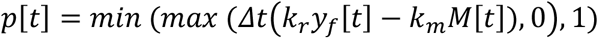

where *k*_*m*_ is a model parameter. As before, bout initiation is subject in addition to the refractory period constraint.

The swim speeds for a given intensity input follow the same equation as that for the initial model:

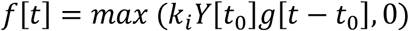

Three special cases of this general model are considered (figures 4A-4C). For variants 2A and 2C, *k*_*f*_ is set to 1, *k*_*b*_ is set to 1 or 0 respectively, and *k*_*i*_ is fitted to data; for variant 2B *k*_*f*_ and *k*_*b*_ are fitted to data and *k*_*i*_ is set to 1.

### Simulation and model fitting

#### Model simulation

Bespoke software was developed in Python for agent-based simulation of the discrete-time model approximation described above. The timestep was 10 ms, the same as the interval between camera frames in the experiments. Each session consisted of 30 s exposure to the simulated moving stimulus, with the final 20 s of behavior analyzed, giving time for a steady state to be reached during the initial 10 s. 30 simulated fish were run for each condition using the same values for the independent variables (height and stimulus speed) as in the two experiments.

#### Cost function

To fit the model to data requires minimizing a cost, here the *relative mean prediction error* for one or more outcome variables, with respect to the model parameters. For example, when considering the single outcome variable of mean swim speed we define the individual prediction error for a given experimental condition (combination of height and stimulus speed) and parameter set as the difference between the mean swim speed observed for that condition and that predicted through model simulation with those parameters. The overall prediction error for an outcome variable is then defined as the root mean square of the individual errors taken over all experimental conditions and the relative prediction error as the overall prediction error divided by the mean value observed for that outcome variable over all experimental conditions. These relative prediction errors are dimensionless and thus no longer sensitive to the subjective choice of units for an outcome variable (e.g., mm/s vs m/s for speed). When fitting for multiple outcome variables simultaneously (usually bout rate and bout intensity) the mean of the relative prediction errors for each variable yields a scalar cost value as required for optimization in a way that balances in an objective way the importance given to each variable.

#### Optimization process

The optimization process was a semi-automatic method based on repeated grid search. At each iteration an n-dimensional rectangular grid of variables was generated and each point of the grid transformed to a set of model parameters. Relative prediction errors were calculated for each parameter set and the optimal set taken as the central point for the grid for the next iteration, resulting in a series of iterations with the grid translated at each step until the best point remained in the center. The pitch of each dimension of the grid was then reduced and the process repeated, eventually closing in on an optimal set of model parameters. Decisions about how quickly to reduce the pitch of a dimension were guided by contour maps and influenced by the sensitivity of the prediction error to changes in that dimension. The maps also helped to avoid settling for a local minimum when it seemed possible that a better point might lie somewhere else within the region currently being explored.

## Supporting information

**S1.**
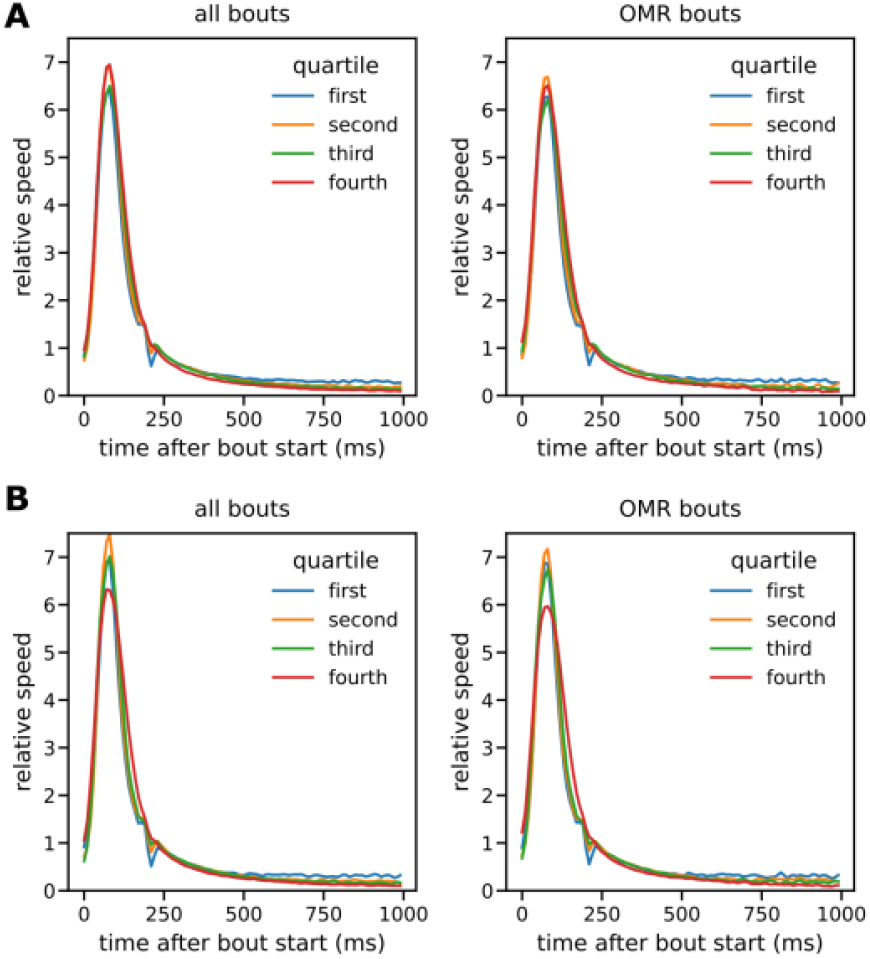
Bout speed profiles. Separate relative speed profiles for bouts split into quartiles by measures of overall bout intensity other than bout initial speed: peak bout speed (A) and total bout displacement (B). The relative speed profiles are again essentially independent of bout intensity whichever measure is used and the profile for OMR-consistent bouts is similar to that for all bouts taken together.

**S2.**
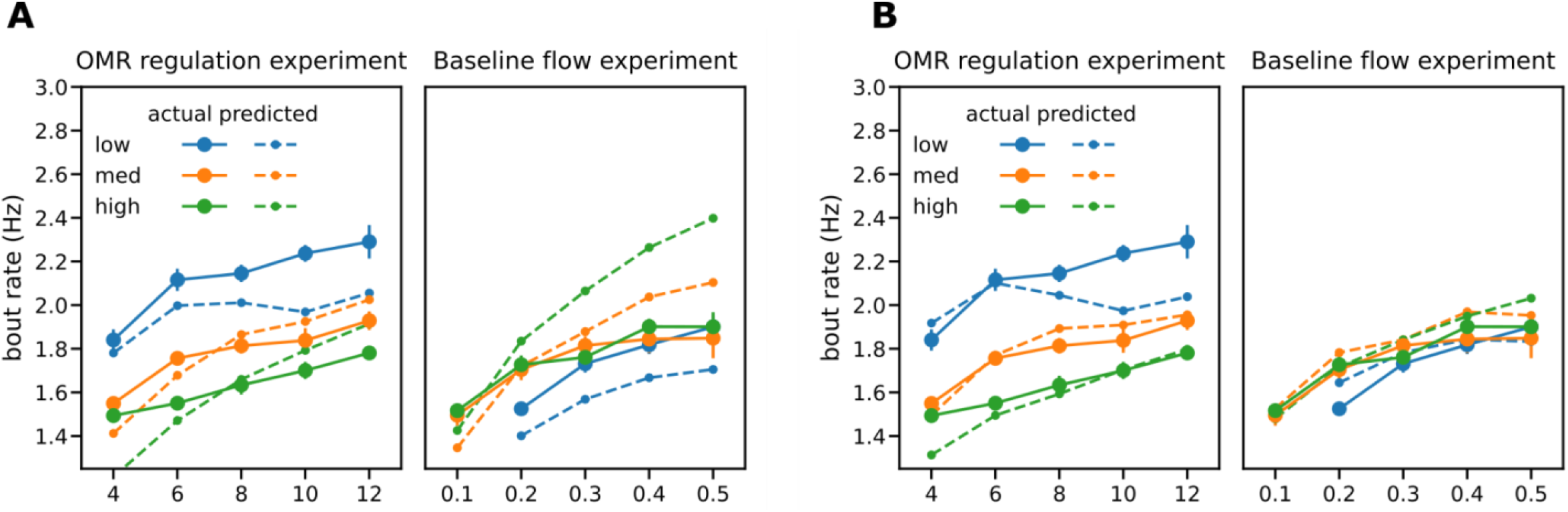
Bout rates predictions for basic and enhanced bout generators. (A-B) Model predictions for bout rates when bout intensity is set to the mean observed values for each experimental condition. (A) Basic bout generator. For the baseline flow procedure, observed bout rates are insensitive to height but the basic model cannot reproduce this finding. (B) Enhanced bout generator with motor inhibition. The qualitative findings can now be reproduced.

**S3.**
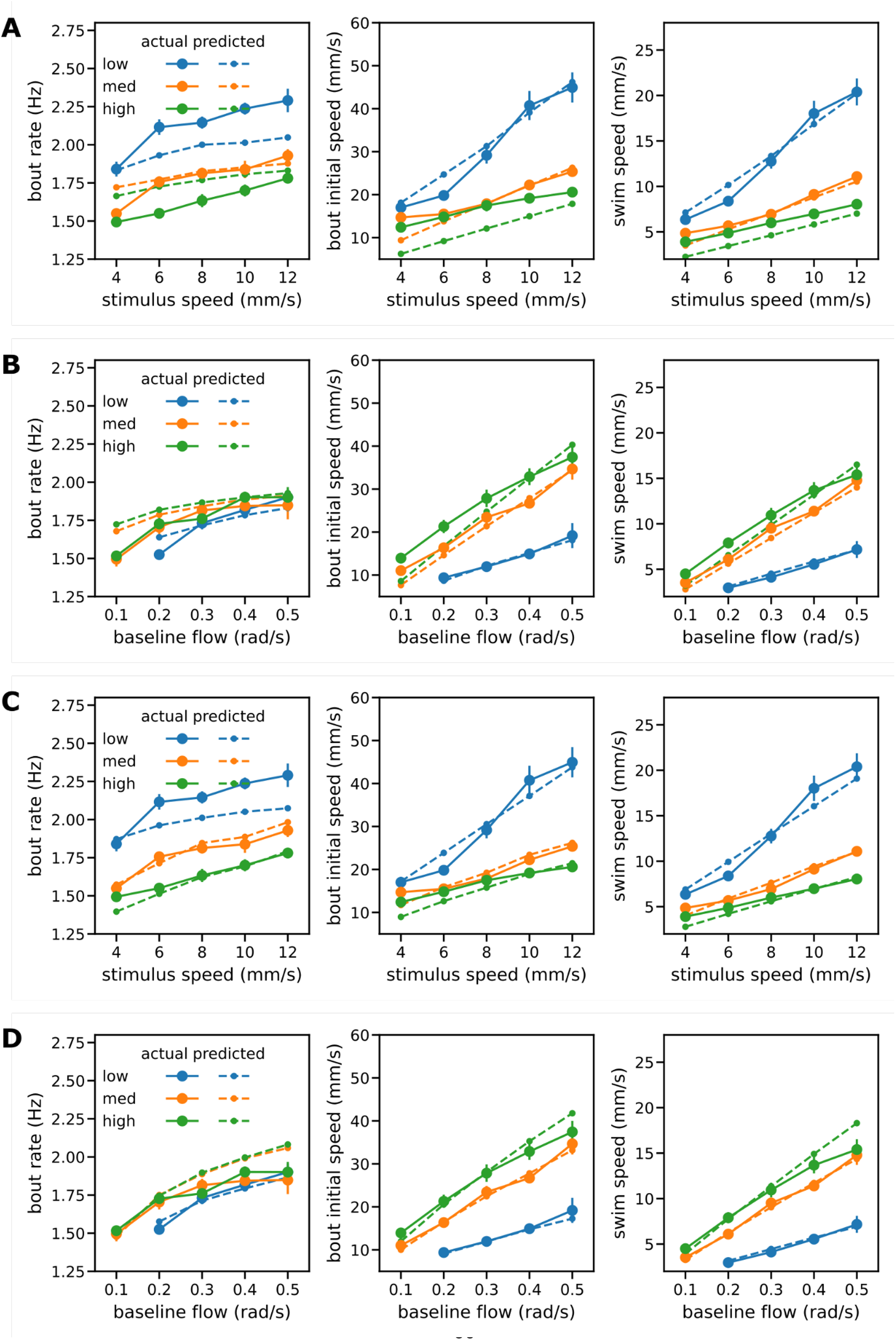
Performance of two-factor model variants A and B. (A-B) Comparison of observed bout rates, bout intensities and swim speeds for the OMR regulation procedure (A) and baseline flow procedure (B) with predictions from model variant A. (C-D) Comparison of observed bout rates, bout intensities and swim speeds for the OMR regulation procedure (C) and baseline flow procedure (D) with predictions from model variant B.

## Bibliography

1. Srinivasan MV. Visual control of navigation in insects and its relevance for robotics. Curr Opin Neurobiol. 2011 Aug 1;21(4):535–43.

2. Collett T, Nalbach HO, Wagner H. Visual stabilization in arthropods. Rev Oculomot Res. 1993;5:239– 63.

3. Arnold GP. Rheotropism in Fishes. Biol Rev. 1974 Nov 1;49(4):515–76.

4. Orger MB, Polavieja GG de. Zebrafish Behavior: Opportunities and Challenges. Annu Rev Neurosci. 2017;40(1):125–47.

5. Kretschmer F, Tariq M, Chatila W, Wu B, Badea TC. Comparison of optomotor and optokinetic reflexes in mice. J Neurophysiol. 2017 Jul 1;118(1):300–16.

6. Warren WH, Kay BA, Zosh WD, Duchon AP, Sahuc S. Optic flow is used to control human walking. Nat Neurosci. 2001 Feb;4(2):213–6.

7. Portugues R, Engert F. The neural basis of visual behaviors in the larval zebrafish. Curr Opin Neurobiol. 2009 Dec 1;19(6):644–7.

8. Ahrens MB, Orger MB, Robson DN, Li JM, Keller PJ. Whole-brain functional imaging at cellular resolution using light-sheet microscopy. Nat Methods. 2013 May;10(5):413–20.

9. Portugues R, Engert F. Adaptive Locomotor Behavior in Larval Zebrafish. Front Syst Neurosci [Internet]. 2011 [cited 2018 Feb 18];5. Available from: https://www.frontiersin.org/articles/10.3389/fnsys.2011.00072/full

10. Ahrens MB, Li JM, Orger MB, Robson DN, Schier AF, Engert F, et al. Brain-wide neuronal dynamics during motor adaptation in zebrafish. Nature. 2012 May;485(7399):471–7.

11. Naumann EA, Fitzgerald JE, Dunn TW, Rihel J, Sompolinsky H, Engert F. From Whole-Brain Data to Functional Circuit Models: The Zebrafish Optomotor Response. Cell. 2016 Nov 3;167(4):947-960.e20.

12. Markov DA, Petrucco L, Kist AM, Portugues R. A cerebellar internal model calibrates a feedback controller involved in sensorimotor control. Nat Commun. 2021 Nov 18;12(1):6694.

13. Bak-Coleman J, Smith D, Coombs S. Going with, then against the flow: evidence against the optomotor hypothesis of fish rheotaxis. Anim Behav. 2015 Sep 1;107:7–17.

14. Kist AM, Portugues R. Optomotor Swimming in Larval Zebrafish Is Driven by Global Whole-Field Visual Motion and Local Light-Dark Transitions. Cell Rep. 2019 Oct 15;29(3):659-670.e3.

15. Krakauer JW, Ghazanfar AA, Gomez-Marin A, MacIver MA, Poeppel D. Neuroscience Needs Behavior: Correcting a Reductionist Bias. Neuron. 2017 Feb 8;93(3):480–90.

16. Orger MB, Kampff AR, Severi KE, Bollmann JH, Engert F. Control of visually guided behavior by distinct populations of spinal projection neurons. Nat Neurosci. 2008 Mar;11(3):327–33.

17. Budick SA, O’Malley DM. Locomotor repertoire of the larval zebrafish: swimming, turning and prey capture. J Exp Biol. 2000 Sep 1;203(17):2565–79.

18. Huang Y-Y, Neuhauss SCF. The optokinetic response in zebrafish and its applications. Front Biosci. 2008;13:1899–916.

19. Koenderink JJ, van Doorn AJ. Facts on optic flow. Biol Cybern. 1987 Jun 1;56(4):247–54.

20. David CT. Compensation for height in the control of groundspeed by Drosophila in a new, ‘barber’s pole’ wind tunnel. J Comp Physiol. 1982 Dec 1;147(4):485–93.

21. Srinivasan MV, Zhang SW, Chahl JS, Barth E, Venkatesh S. How honeybees make grazing landings on flat surfaces. Biol Cybern. 2000 Aug 1;83(3):171–83.

22. Kawashima T, Zwart MF, Yang C-T, Mensh BD, Ahrens MB. The Serotonergic System Tracks the Outcomes of Actions to Mediate Short-Term Motor Learning. Cell. 2016 Nov 3;167(4):933-946.e20.

23. Neuhauss SCF, Biehlmaier O, Seeliger MW, Das T, Kohler K, Harris WA, et al. Genetic Disorders of Vision Revealed by a Behavioral Screen of 400 Essential Loci in Zebrafish. J Neurosci. 1999 Oct 1;19(19):8603–15.

24. Severi KE, Portugues R, Marques JC, O’Malley DM, Orger MB, Engert F. Neural Control and Modulation of Swimming Speed in the Larval Zebrafish. Neuron. 2014 Aug 6;83(3):692–707.

25. Maaswinkel H, Li L. Spatio-temporal frequency characteristics of the optomotor response in zebrafish. Vision Res. 2003 Jan 1;43(1):21–30.

26. Engert F. Fish in the matrix: motor learning in a virtual world. Front Neural Circuits [Internet]. 2013 [cited 2018 Feb 27];6. Available from: https://www.frontiersin.org/articles/10.3389/fncir.2012.00125/full

27. Merton PA. Speculations on the servo-control of movement. In: Wolstenholme GEW, editor. The Spinal Cord. London: Churchill; 1953. p. 247–55.

28. Åström KJ, Murray R. Feedback Systems: An Introduction for Scientists and Engineers. 2nd ed. Princeton University Press; 2021.

29. Dayan P, Abbott LF. Theoretical Neuroscience: Computational and Mathematical Modelling of Neural Systems. Cambridge, MA, USA: MIT Press; 2001.

30. Gerstner W, Kistler WM, Naud R, Paninski L. Neuronal Dynamics: From Single Neurons to Networks and Models of Cognition [Internet]. Cambridge University Press; 2014. 591 p. Available from: https://neuronaldynamics.epfl.ch/online/index.html

31. Portugues R, Haesemeyer M, Blum ML, Engert F. Whole-field visual motion drives swimming in larval zebrafish via a stochastic process. J Exp Biol. 2015 May 1;218(9):1433–43.

32. Kubo F, Hablitzel B, Dal Maschio M, Driever W, Baier H, Arrenberg AB. Functional Architecture of an Optic Flow-Responsive Area that Drives Horizontal Eye Movements in Zebrafish. Neuron. 2014 Mar 19;81(6):1344–59.

33. Wang K, Hinz J, Haikala V, Reiff DF, Arrenberg AB. Selective processing of all rotational and translational optic flow directions in the zebrafish pretectum and tectum. BMC Biol. 2019 Mar 29;17(1):29.

34. Miller CS, Johnson DH, Schroeter JP, Myint LL, Glantz RM. Visual Signals in an Optomotor Reflex: Systems and Information Theoretic Analysis. J Comput Neurosci. 2002 Jul 1;13(1):5–21.

35. Tresilian J. Sensorimotor Control and Learning. Palgreave Macmillan; 2012.

36. Moltz H. Contemporary instinct theory and the fixed action pattern. Psychol Rev. 1965;72(1):27–47.

37. Barlow GW. Modal action patterns. In: Sebeok TA, editor. How animals communicate [Internet]. Bloomington, IN: Indiana University Press; 1977. p. 98–134. Available from: https://publish.iupress.indiana.edu/read/how-animals-communicate/section/6f9a6531-694b-4058-94b0-8b0819851348#ch6

38. Marr D. Vision: A Computational Investigation into the Human Representation and Processing of Visual Information. MIT Press; 2010. 429 p.

39. Wiggin TD, Anderson TM, Eian J, Peck JH, Masino MA. Episodic swimming in the larval zebrafish is generated by a spatially distributed spinal network with modular functional organization. J Neurophysiol. 2012 Aug 1;108(3):925–34.

40. Olive R, Wolf S, Dubreuil A, Bormuth V, Debrégeas G, Candelier R. Rheotaxis of Larval Zebrafish: Behavioral Study of a Multi-Sensory Process. Front Syst Neurosci [Internet]. 2016 [cited 2022 Jan 31];10. Available from: https://www.frontiersin.org/article/10.3389/fnsys.2016.00014

41. Imada H, Hoki M, Suehiro Y, Okuyama T, Kurabayashi D, Shimada A, et al. Coordinated and Cohesive Movement of Two Small Conspecific Fish Induced by Eliciting a Simultaneous Optomotor Response. PLOS ONE. 2010 Jun 22;5(6):e11248.

42. Srinivasan MV, Bernard GD. The pursuit response of the housefly and its interaction with the optomotor response. J Comp Physiol. 1977 Jan 1;115(1):101–17.

43. Reback J, jbrockmendel, McKinney W, Bossche JV den, Augspurger T, Cloud P, et al. pandas-dev/pandas: Pandas 1.2.5 [Internet]. Zenodo; 2021 [cited 2021 Nov 14]. Available from: https://zenodo.org/record/5013202

44. R Core Team. R: A Language and Environment for Statistical Computing. [Internet]. Vienna, Austria: R Foundation for Statistical Computing; 2021. Available from: https://www.R-project.org/

